# T cell response following anti COVID-19 BNT162b2 vaccination is maintained against the SARS-CoV-2 Omicron B.1.1.529 variant of concern

**DOI:** 10.1101/2022.01.19.476497

**Authors:** Hila Cohen, Shahar Rotem, Uri Elia, Gal Bilinsky, Itzchak Levy, Theodor Chitlaru, Erez Bar-Haim

## Abstract

The progression of the COVID-19 pandemic leads to the emergence of variants of concern (VOC), which may compromise the efficacy of the currently administered vaccines. Antigenic drift can potentially bring about a reduced protective T cell immunity and consequently to more severe disease manifestations. To assess this possibility, the T cell responses to the wild-type, Wuhan-1 SARS-CoV-2 ancestral spike protein and Omicron B.1.1.529 spike protein were compared. Accordingly, peripheral blood mononuclear cells (PBMC) were collected from 8 healthy volunteers 4-5 months following a third vaccination with BNT162b2, and stimulated with overlapping peptide libraries representing the spike of either the ancestral or Omicron SARS-CoV- 2 virus variants. Quantification of the specific T cells was carried out by a fluorescent ELISPOT assay, monitoring interferon-gamma (IFNg), interleukin-10 (IL-10) and interleukin-4 (IL-4) secreting cells. For all the examined individuals, comparable level of reactivity to both forms of spike protein were determined. In addition, a dominant Th1 response was observed, manifested mainly by IFNg secreting cells and only limited numbers of IL-10 and IL-4 secreting cells. The data demonstrates a stable T cell activity to the emerging Omicron variant in the tested individuals, therefore the protective immunity to the variant following BNT162b2 vaccination is not significantly affected.

## Introduction

Omicron B.1.1.529 is currently the prevalent variant of concern (VOC) amongst emerging SARS-CoV-2 variants [1]. The Omicron variant was first described in November 2021, and since then is rapidly spreading worldwide [1,2]. It bears 26-32 mutations in the spike protein compared to the Wuhan-1 SARS-CoV-2 sequence [1], many of these mutations being located in the receptor binding domain (RBD). As was shown for other VOCs [3], mutations in the neutralizing sites of the spike protein weaken the neutralizing potential of antibodies and consequently may lead to enhanced immunological escape. This aspect has tremendous public health implications considering the massive on-going vaccination world-wide campaigns based on the antigenic specificity of the primordial SARS-CoV-2 strain.

As of today, precise correlates of protection against SARS-CoV-2 are not fully defined. It is clear that neutralizing antibody response is essential for blocking viral attachment and entry to host cells, and that T cells are playing a central role in diminishing viral spread in the host, therefore alleviating the severity of the disease manifestation [4]. Accordingly, for several emerging VOCs, it was shown that a lower neutralizing antibody response is correlated with lower efficiency of the vaccine and higher levels of immune-breakthrough infections [4,5]. Considering the antibody titer kinetics following vaccination and their potential waning below the neutralizing levels, it is essential to maintain protective T cell memory responses, which are expected to exhibit significant longevity [6–8]. Immune escape from humoral response is mostly a result of specific mutations of a given antigen, which occur in a convergent microevolutionary process and therefore affects equally different individuals. Conversely, T cell response have a divergent character, distinctly affecting various individuals due to HLA polymorphism, and therefore unique mutations in immunodominant epitopes are less likely to affect the T cell responses globally. Weakening of T cell immunity against VOC may occur as a consequence of antigenic drift that leads to accumulated mutations underlying immunity [9].

T cell responses may provide protection from SARS-CoV-2 even in the absence of antibody response [10]. Specifically, high levels IFNg secreting cells responsive to antigenic stimulation with the SARS-CoV-2 spike protein correlate with less severe COVID-19 disease manifestations [10]. Monitoring T cell response is experimentally more challenging than quantification of humoral responses, requiring availability of viable cells responding to antigen stimulation. Most studies characterize T cell responses by activation-induced marker (AIM) elevation and cytokine intracellular staining, monitored by flow cytometry and by ELISPOT assays [11]. In the current study we determined the level of the T cell reactivity in response to the ancestral Wuhan-1 SARS-CoV-2 spike and Omicron B.1.1.529 variant spike in healthy individuals immunized with the BNT162b2 vaccine. The study evidenced a similar, dominant Th1 response to both versions of the spike protein suggesting that stable T cell immunity is maintained against the currently prevalent Omicron variant.

## Materials and Methods

### PBMC Isolation

Blood was collected from 8 healthy individuals, within 4-5 months from a third BNT162b2 vaccine. Blood was collected into sodium-heparin tubes (vacutainer, BD) and processed within 2 hours of collection. Peripheral blood mononuclear cells (PBMC) were isolated by density gradient sedimentation using Ficoll-Paque (Sigma-Aldrich) according to the manufacturer’s protocol, cells were then washed once in PBS and immediately processed for ELISPOT assay.

### ELISPOT Assay

Three color Fluorescent ELISPOT assay (FluoroSpot) was performed in strict adherence to the manufacturer’s protocol (Human IFN-γ/IL-4/IL-10 Three-Color FluoroSpot, ImmunoSpot, Cleveland, OH, USA). PBMC were resuspended in FCS-free, CTL-Test media (ImmunoSpot) and plated in a 96 well PVDF membrane plates, 3×105 cells/well. Cells were either left unstimulated, stimulated with SARS-CoV-2 spike protein overlapping peptide library or stimulated with 5μg/ml of Phytohaemagglutinin (PHA) (Sigma-Aldrich) as a positive control. Commercially available peptide pools (15mer sequences with 11 amino acids overlap) covering the full length of the Wuhan-1 SARS-CoV-2 (wild type) or Omicron B.1.1.529 variant spike (peptides & elephants GmbH, Hennigsdorf, Germany) were used for PBMC stimulation. Peptide pools were dissolved in DMSO and used in a final concentration of 200 μg/ml (0.6 μg/ml per peptide), DMSO final concentration was below 0.1%. The plate layout is presented in Supplementary Figure S1. PBMC were stimulated for 48 hours, and the frequency of cytokine-secreting cells was quantified with ImmunoSpot S6 Ultimate reader with the 520, 600 and 690 nm filters to allow enumeration of IFNg, IL-10 and IL-4 expressing cells, respectively. Data were analyzed with the ImmunoSpot software version 7.0.30.2 (ImmunoSpot). Positive spots overlapped by the 520 nm and 600 or 690 nm filters were considered as potential artefacts and excluded.

### Institutional Review Board Statement

All subjects formally informed their explicit consent of participation prior to their inclusion in the study. The study was conducted in accordance with the Declaration of Helsinki, and the protocol was approved by the Ethics Committee of the Sheba Medical Center (SMC-20-7026).

## Results and Discussion

Cellular immunity is instrumental in preventing severe COVID-19. In the current study which included eight healthy individuals (referred as donors), vaccinated in Israel 3 times with the mRNA BNT162b2 vaccine, we sought to determine and compare the level and type of T cell response to either Wuhan-1 SARS-CoV-2 spike or Omicron B.1.1.529 variant. The donors were 20- 52 years old (average 27.1,) and included 3 males and 5 females. No COVID-19 history was documented for any of the donors; furthermore, considering the accurate epidemiologic registering, customary in Israel, it is conceivable that they were not previously infected.

PBMC collected from the donors were stimulated with a mixture of 315 peptides, 15 amino-acids long, spanning the entire spike protein. The study inspected both the type and the level of response, as determined by the number of cytokine-secreting cells in a fluorescent ELISPOT assay. Following antigenic stimulation with the spike-derived peptides, a predominant IFNg response was observed in all the examined individuals, ranging from 50 to 400 secreting cells per 10^6^ PBMC, and a lower IL-10 response, ranging from 15 to 82 cells per 10^6^ PBMC (figure 1 and 1S). The experimental setup included a 48 hours antigen stimulation step, to allow manifestation of reactivity of IL-4-expressing T cell clones potentially present in the samples, however, almost undetectable levels of IL-4 reactivity was determined, in accordance with previous data pertaining to the responses elicited by the BNT162b2 and m1273 vaccines [2,4,8–13].Comparison of the average response to the wild type and Omicron spike (Figure 2), indicated only a slight, non-significant decrease, from 201 IFNg-secreting cells following activation with the wild type spike, to 188 cells responding to the Omicron spike. The IFNg response was higher than that of IL-10, the average ratio of IFNg/IL-10 response being 4.9, indicating a dominant Th1 response with no significant Th2 response.

**Figure 1.**
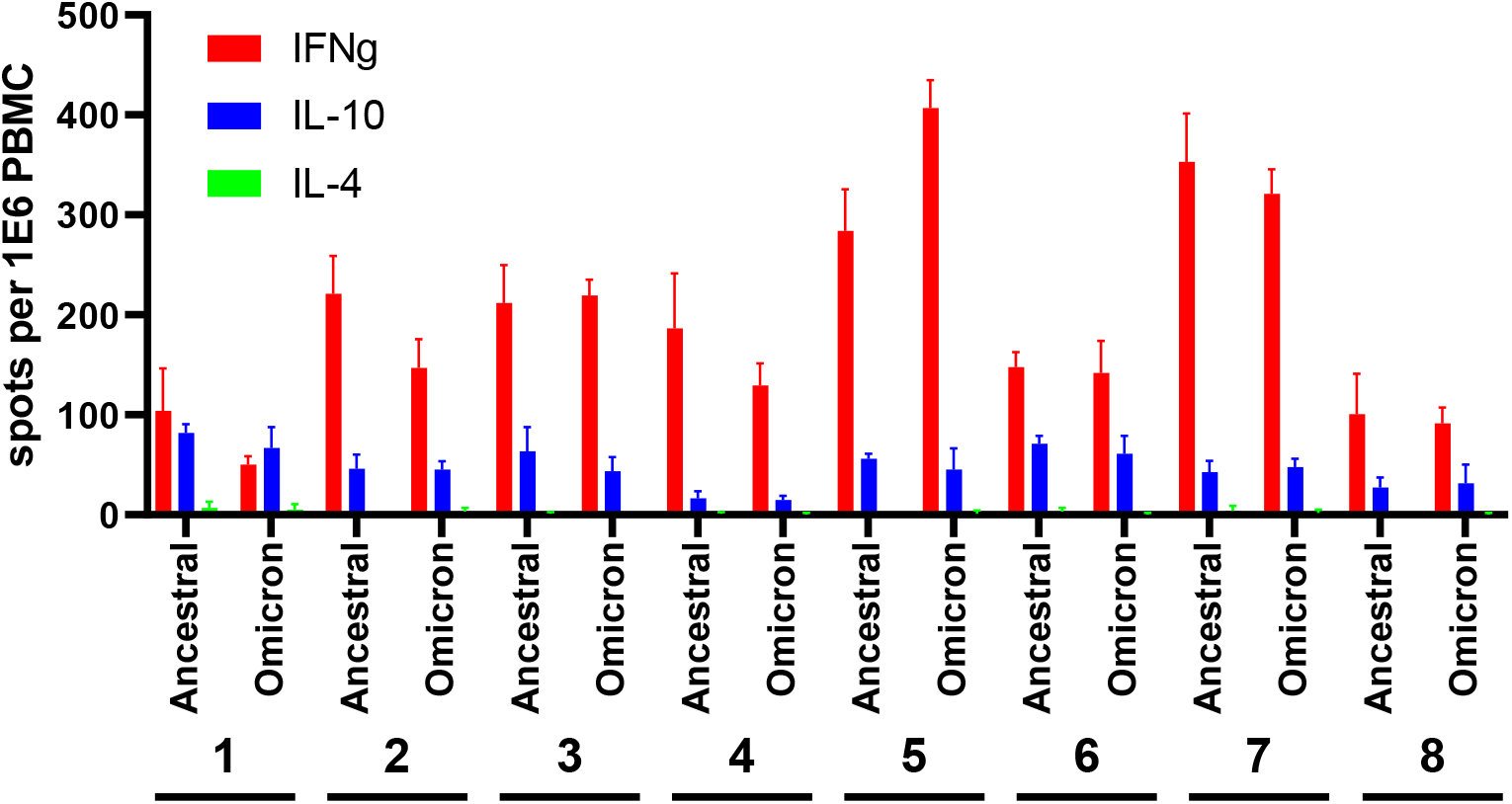
T cell response to the ancestral and Omicron SARS-CoV-2 spike in BNT162b2-vaccinated individuals. PBMCs were stimulated with ancestral or Omicron spike-derived overlapping peptides. IFNg, IL-10 and IL-4-secreting cells were quantified in a FluoroSpot assay. Data represent the average and standard deviation of four replications of each experimental group.

**Figure 2.**
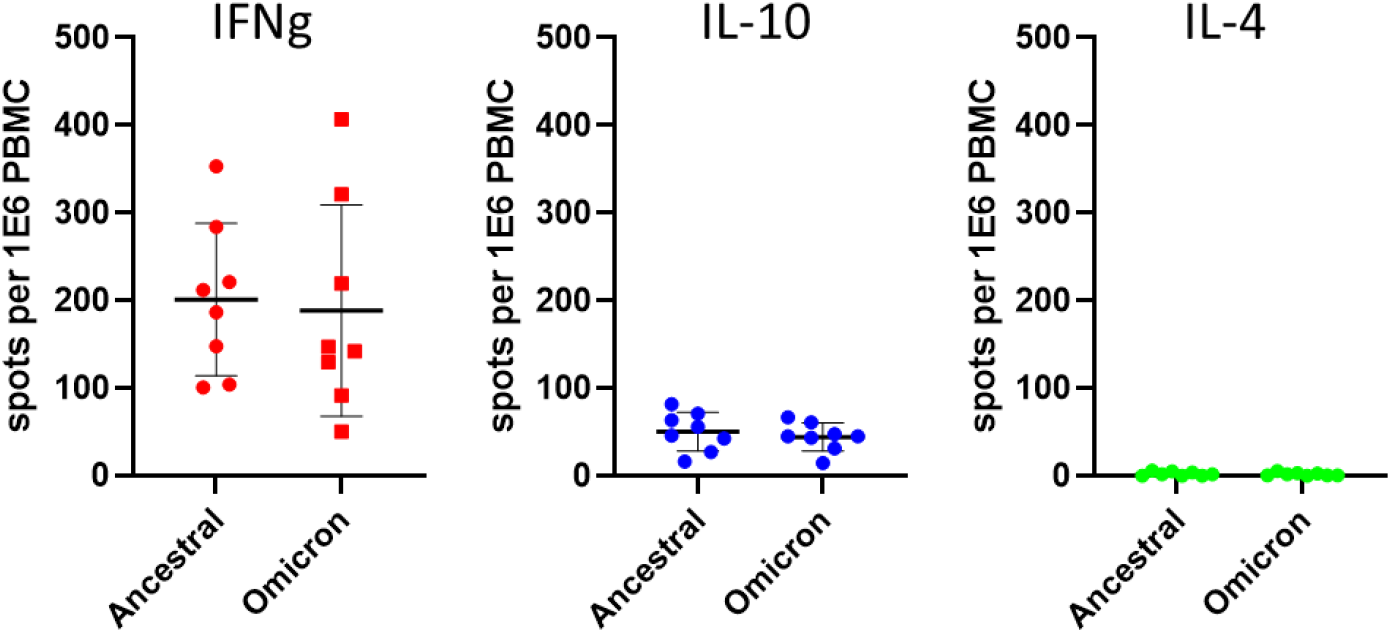
Comparative analysis of the T cell response to ancestral and Omicron spike protein. PBMCs were stimulated as described in Figure 1. Each dot represents the average of four measurements of the same sample. Average and standard deviation of the data from the independent donors for each cytokine and antigen are presented.

The essentiality of T cells for protection against COVID-19 is well documented [2,4,10], therefore confirmation of T cell reactivity towards emerging VOCs is of outmost importance. As newer emerging VOCs are identified, maintenance of long-term protective immunity of vaccinated individuals represents a public health concern of high priority. It was suggested that in the case of several VOCs, convalescent and vaccinated individuals exhibited some escape from humoral immunity, while revealing normal uncompromised T cell reactivity [5, 14].

In the present study, by analyzing the response in individuals following 3 BNT162b2vaccines, we show a dominant Th1 response to the Omicron variant spike protein, which correlates with protective immunity [6]. The T cell response to both the ancestral and Omicron was of commensurate levels (figure 2) in line with several recent reports [2,12,13]. Since our data show comparable level of response to both ancestral and Omicron spike, it is reasonable to estimate that the CD4 and CD8 composition of the T cell compartment remains steady for the response to both variants. The added value of a third vaccination for achieving Omicron antibody neutralization was previously demonstrated [15], and future studies will address the relevance of the third vaccine administration in maintenance of the T cells responses as well. In addition, additional studies are expected to monitor other population fractions, especially older people at high risk of developing severe forms of COVID-19.

## Supporting information

Figure S1: FluoroSpot plate layout and quantitation

## Supplementary Materials

The following supporting information are available: Figure S1: FluoroSpot plate layout and quantitation.

## Author Contributions

Conceptualization and investigation: HC, SR, UE, GB, IL and EBH; assessment of data and writing: TC and EBH; EBH supervised the project.

## Funding

This study was funded by an intra-mural grant of the Israel Institute for Biological Research.

## Acknowledgments

We are grateful to Dr. Emanuelle Mamroud for productive discussions and for her enthusiastic support. We would like to thank Ms. Yael Shlomo for excellent administrative support.

## Conflicts of Interest

The authors declare no conflict of interest.

## Notes

### Competing Interest Statement

The authors have declared no competing interest.

## References

1. Pulliam, J.R.C.; van Schalkwyk, C.; Govender, N.; von Gottberg, A.; Cohen, C.; Groome, M.J.; Dushoff, J.; Mlisana, K.; Moultrie, H. Increased risk of SARS-CoV-2 reinfection associated with emergence of the Omicron variant in South Africa. medRxiv 2021, 2021.2011.2011.21266068, doi:10.1101/2021.11.11.21266068.

2. Keeton, R.; Tincho, M.B.; Ngomti, A.; Baguma, R.; Benede, N.; Suzuki, A.; Khan, K.; Cele, S.; Bernstein, M.; Karim, F.; et al. SARS-CoV-2 spike T cell responses induced upon vaccination or infection remain robust against Omicron. medRxiv 2021, 2021.2012.2026.21268380, doi:10.1101/2021.12.26.21268380.

3. Liu, C.; Ginn, H.M.; Dejnirattisai, W.; Supasa, P.; Wang, B.; Tuekprakhon, A.; Nutalai, R.; Zhou, D.; Mentzer, A.J.; Zhao, Y.; et al. Reduced neutralization of SARS-CoV-2 B.1.617 by vaccine and convalescent serum. Cell 2021, 184, 4220–4236.e4213, doi:https://doi.org/10.1016/j.cell.2021.06.020.

4. Cevik, M.; Grubaugh, N.D.; Iwasaki, A.; Openshaw, P. COVID-19 vaccines: Keeping pace with SARS-CoV-2 variants. Cell 2021, 184, 5077–5081, doi:10.1016/j.cell.2021.09.010.

5. Kustin, T.; Harel, N.; Finkel, U.; Perchik, S.; Harari, S.; Tahor, M.; Caspi, I.; Levy, R.; Leshchinsky, M.; Ken Dror, S.; et al. Evidence for increased breakthrough rates of SARS-CoV-2 variants of concern in BNT162b2-mRNA-vaccinated individuals. Nature Medicine 2021, 27, 1379–1384, doi:10.1038/s41591-021-01413-7.

6. Altmann, D.M.; Reynolds, C.J.; Boyton, R.J. SARS-CoV-2 variants: Subversion of antibody response and predicted impact on T cell recognition. Cell Reports Medicine 2021, 2, 100286, doi:https://doi.org/10.1016/j.xcrm.2021.100286.

7. Baraniuk, C. How long does covid-19 immunity last? BMJ 2021, 373, n1605, doi:10.1136/bmj.n1605.

8. Reynolds, C.J.; Gibbons, J.M.; Pade, C.; Lin, K.-M.; Sandoval, D.M.; Pieper, F.; Butler, D.K.; Liu, S.; Otter, A.D.; Joy, G.; et al. Heterologous infection and vaccination shapes immunity against SARS-CoV-2 variants. Science 0, eabm0811, doi:doi:10.1126/science.abm0811.

9. Altmann, D.M.; Boyton, R.J.; Beale, R. Immunity to SARS-CoV-2 variants of concern. Science 2021, 371, 1103–1104, doi:doi:10.1126/science.abg7404.

10. Noh, J.Y.; Jeong, H.W.; Kim, J.H.; Shin, E.-C. T cell-oriented strategies for controlling the COVID-19 pandemic. Nature Reviews Immunology 2021, 21, 687–688, doi:10.1038/s41577-021-00625-9.

11. Ogbe, A.; Kronsteiner, B.; Skelly, D.T.; Pace, M.; Brown, A.; Adland, E.; Adair, K.; Akhter, H.D.; Ali, M.; Ali, S.-E.; et al. T cell assays differentiate clinical and subclinical SARS-CoV-2 infections from cross-reactive antiviral responses. Nature Communications 2021, 12, 2055, doi:10.1038/s41467-021-21856-3.

12. GeurtsvanKessel, C.H.; Geers, D.; Schmitz, K.S.; Mykytyn, A.Z.; Lamers, M.M.; Bogers, S.; Gommers, L.; Sablerolles, R.S.G.; Nieuwkoop, N.N.; Rijsbergen, L.C.; et al. Divergent SARS CoV-2 Omicron-specific T- and B-cell responses in COVID-19 vaccine recipients. medRxiv 2021, 2021.2012.2027.21268416, doi:10.1101/2021.12.27.21268416.

13. Naranbhai, V.; Nathan, A.; Kaseke, C.; Berrios, C.; Khatri, A.; Choi, S.; Getz, M.A.; Tano-Menka, R.; Ofoman, O.; Gayton, A.; et al. T cell reactivity to the SARS-CoV-2 Omicron variant is preserved in most but not all prior infected and vaccinated individuals. medRxiv 2022, 2022.2001.2004.21268586, doi:10.1101/2022.01.04.21268586.

14. Geers, D.; Shamier, M.C.; Bogers, S.; Hartog, G.d.; Gommers, L.; Nieuwkoop, N.N.; Schmitz, K.S.; Rijsbergen, L.C.; Osch, J.A.T.v.; Dijkhuizen, E.; et al. SARS-CoV-2 variants of concern partially escape humoral but not T cell responses in COVID-19 convalescent donors and vaccine recipients. Science Immunology 2021, 6, eabj1750, doi:doi:10.1126/sciimmunol.abj1750.

15. Nemet, I.; Kliker, L.; Lustig, Y.; Zuckerman, N.; Erster, O.; Cohen, C.; Kreiss, Y.; Alroy-Preis, S.; Regev-Yochay, G.; Mendelson, E.; et al. Third BNT162b2 Vaccination Neutralization of SARS-CoV-2 Omicron Infection. New England Journal of Medicine 2021, doi:10.1056/NEJMc2119358.

