## Supplementary material for "T cell response following anti COVID-19 BNT162b2 vaccination is maintained against the SARS-CoV-2 Omicron B.1.1.529 variant of concern": Figure S1: FluoroSpot plate layout and quantitation

A

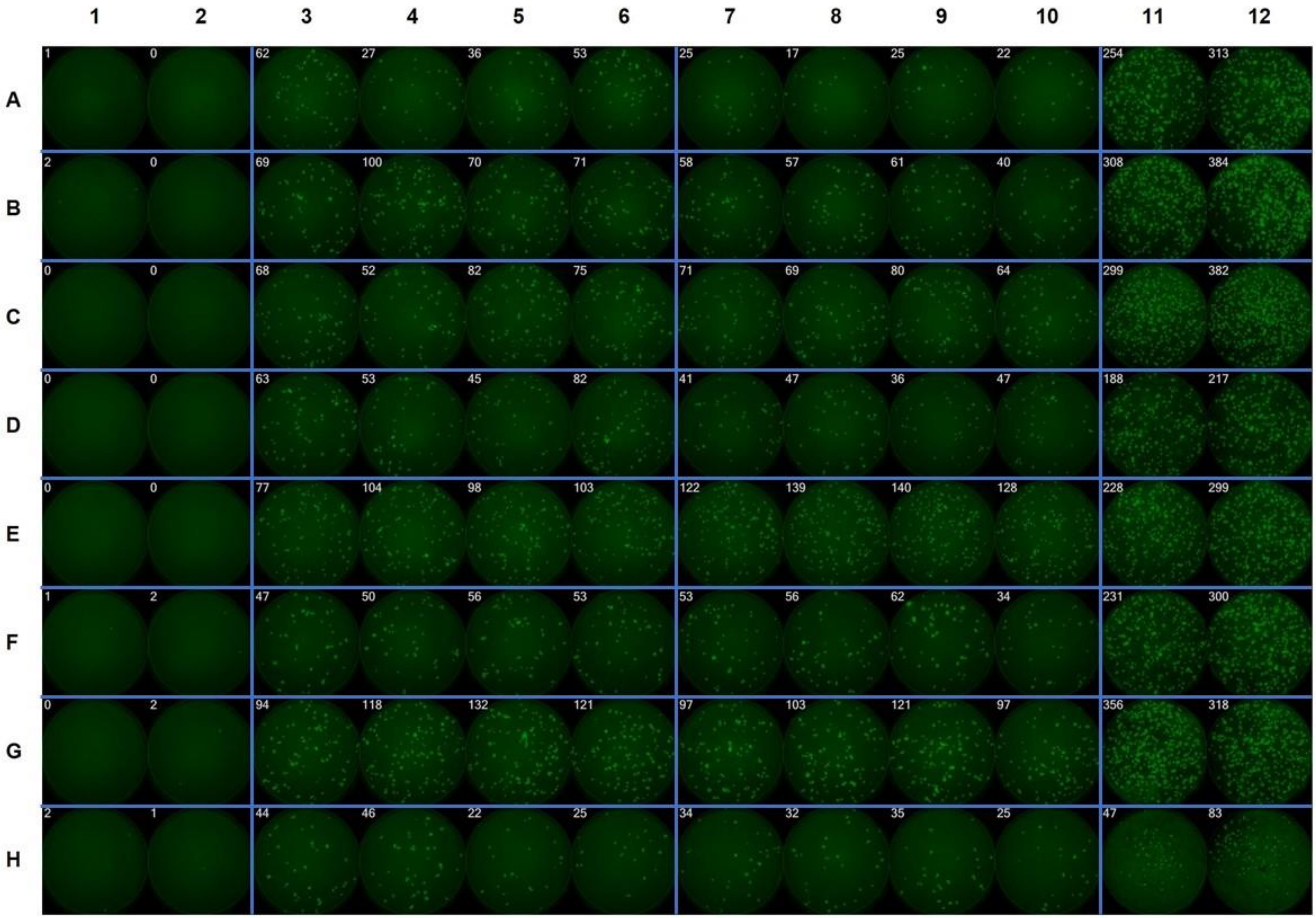

1 2 3 4 5 6 7 8 9 10 11 12
A
1 0 62 27 36 53 25 17 25 22 254 313
B
2 0 69 100 70 71 58 57 61 40 308 384
C
0 0 68 52 82 75 71 69 80 64 299 382
D
0 0 63 53 45 82 41 47 36 47 188 217
E
0 0 77 104 98 103 122 139 140 128 228 299
F
1 2 47 50 56 53 53 56 62 34 231 300
G
0 2 94 118 132 121 97 103 121 97 356 318
H
2 1 44 46 22 25 34 32 35 25 47 83

B

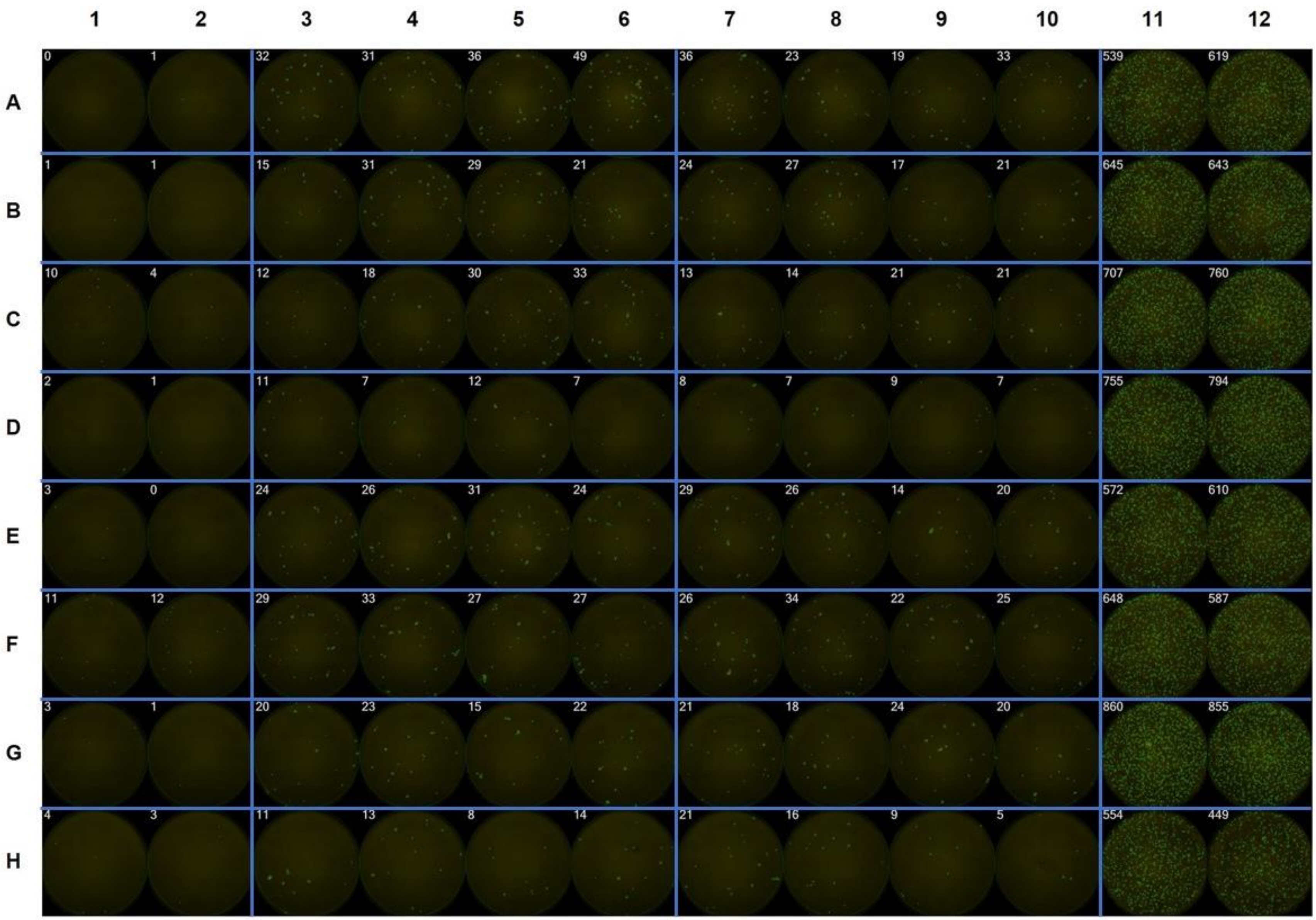
1 2 3 4 5 6 7 8 9 10 11 12
A 0 1 32 31 36 49 36 23 19 33 539 619
B 1 1 15 31 29 21 24 27 17 21 645 643
C 10 4 12 18 30 33 13 14 21 21 707 760
D 2 1 11 7 12 7 8 7 9 7 755 794
E 3 0 24 26 31 24 29 26 14 20 572 610
F 11 12 29 33 27 27 26 34 22 25 648 587
G 3 1 20 23 15 22 21 18 24 20 860 855
H 4 3 11 13 8 14 21 16 9 5 554 449

C

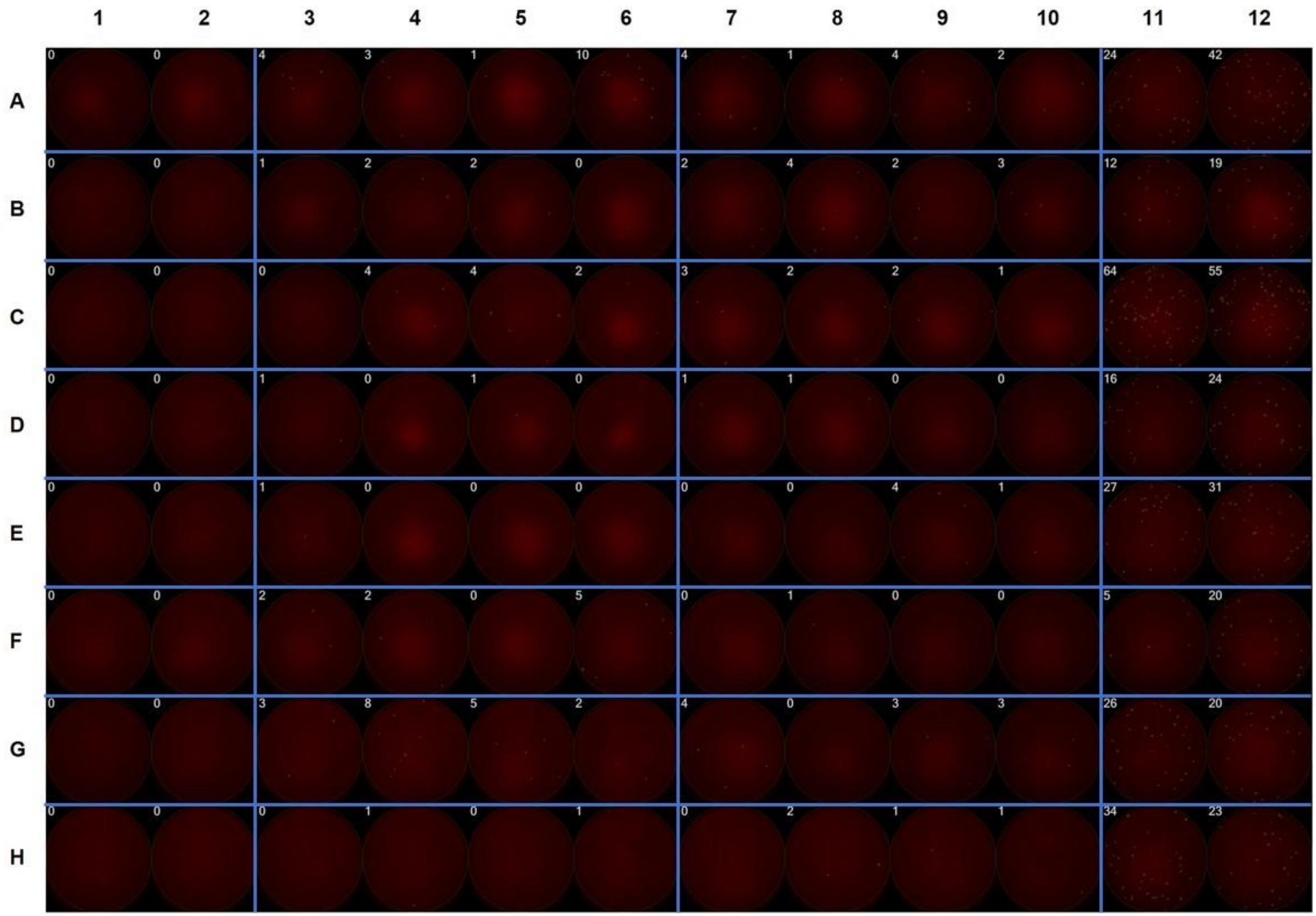


**Figure S1**: FluoroSpot plate layout and quantitation. PBMC ($3x10^5$ per well) were left unstimulated (columns 1, 2), stimulated with ancestral spike (lanes 3-6) or Omicron spike (lanes 7-10) overlapping peptide library pool, or stimulated with L-PHA (5 μg/ml). IFNg (A), IL-10 (B) and IL-4 (C) producing cells were quantified and presented as spots per well. Each raw (A-H) represent one individual donor (Donors 1-8).
